## Supplemental Figures for "Distinct neurochemical influences on fMRI response polarity in the striatum"

**Supplementary Figures**

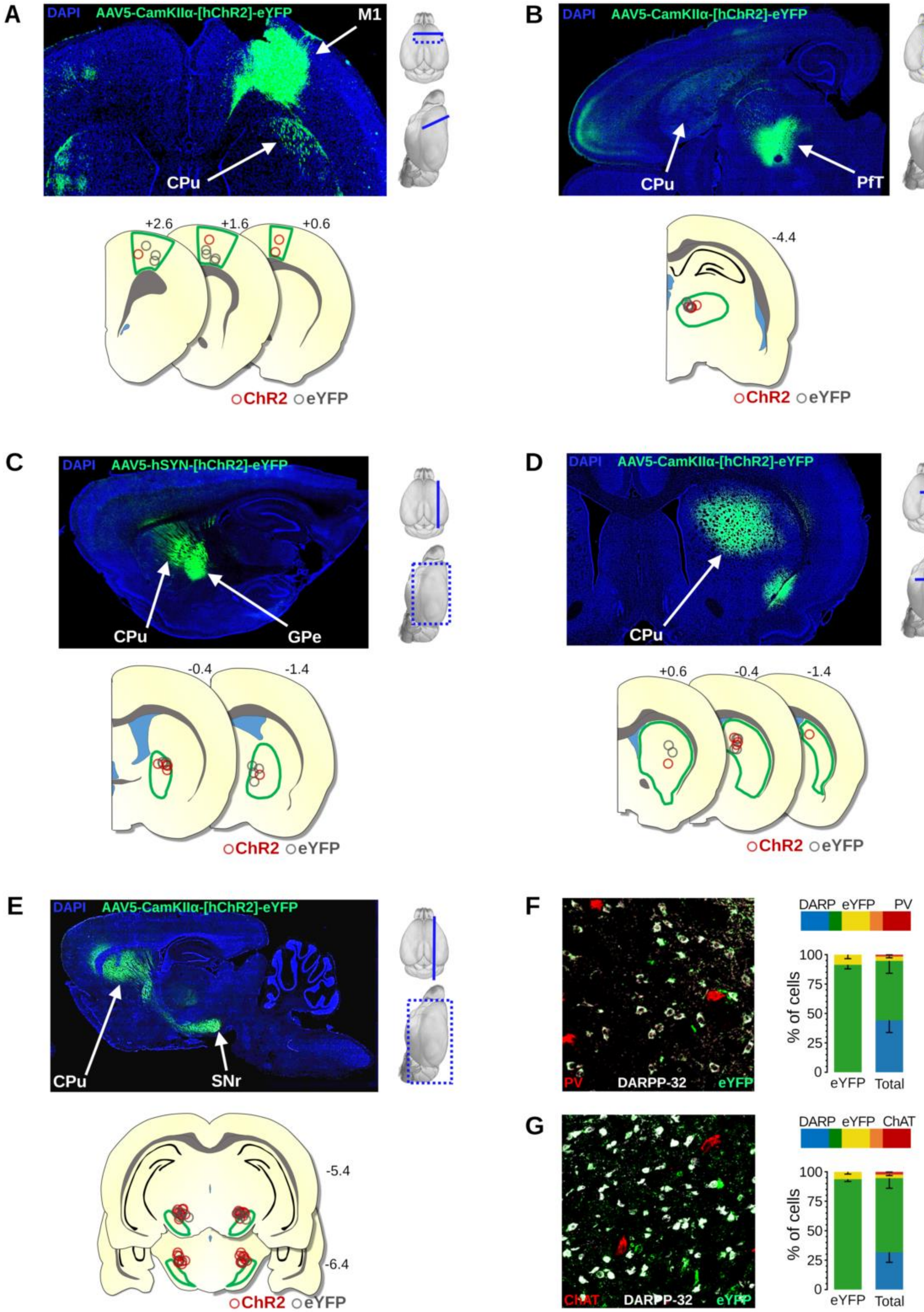
A
DAPI AAV5-CamKIIα-[hChR2]-eYFP
M1
CPu
+2.6
+1.6
+0.6
ChR2 eYFP
B
DAPI AAV5-CamKIIα-[hChR2]-eYFP
CPu
PfT
-4.4
ChR2 eYFP
C
DAPI AAV5-hSYN-[hChR2]-eYFP
CPu
GPe
-0.4
-1.4
ChR2 eYFP
D
DAPI AAV5-CamKIIα-[hChR2]-eYFP
CPu
+0.6
-0.4
-1.4
ChR2 eYFP
E
DAPI AAV5-CamKIIα-[hChR2]-eYFP
CPu
SNr
-5.4
-6.4
ChR2 eYFP
F
PV
DARPP-32
eYFP
DARP eYFP PV
% of cells
100
75
50
25
0
eYFP Total
G
CHAT
DARPP-32
eYFP
DARP eYFP ChAT
% of cells
100
75
50
25
0
eYFP Total

3

**Fig. S1. Confirmation of optogenetic fMRI stimulation targets in Figure 1.** (**A-E**, top) Representative cross-section indicating spread of eYFP for each stimulation target, with 3-D models of the brain to the right, indicating the location of the histological slice. (**A-E**, bottom). Locations of optical fiber tips relative to the anatomical stimulation target ROI outlined in green. Stimulation targets and areas of interest are as follows: Viral expression in M1 injection sites and its spread to CPu. (**B**) Viral expression at the PfT injection site and its spread to CPu. (**C**) Viral expression at the GPe injection site and its spread to CPu. **(D)** Viral expression in the CPu for direct MSN stimulation. (**E**) Viral expression at the CPu injection site and its spread to SNr stimulation sites. (**F-G**) High-resolution 20x magnification image and immunohistochemical verification of DARPP-32 (DARP), eYFP, and either PV (F) or ChAT (G) co-expression in striatum (PV n = 2 brains, 7 slices, 6195 cells; ChAT n = 3 brains, 11 slices, 14181 cells).

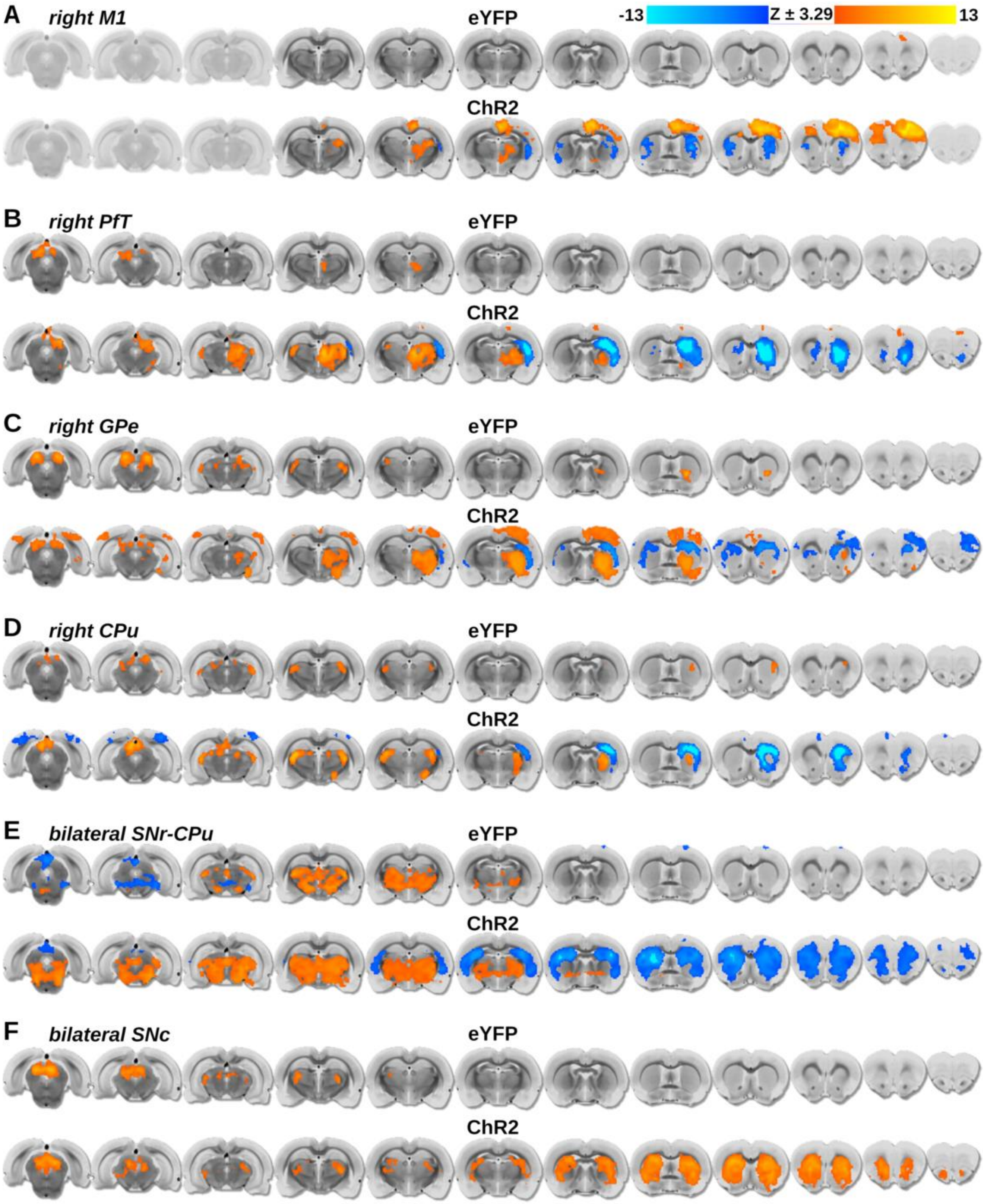


**Fig. S2. Extended 12-slice optogenetic CBV-fMRI response maps acquired during CPu circuit manipulations in Figure 1.** (**A-F**) Response maps acquired from eYFP control and ChR2 subjects following optogenetic stimulation of the circuit indicated at the top left of each set of maps**.** Left-to-right corresponds to 1 mm steps in the posterior-to-anterior direction, with the 5th slice from the right located at the anterior commissure (approximately –0.36 mm AP). Response maps thresholded to $p<0.001$, FWE corrected to $\alpha < 0.01$.

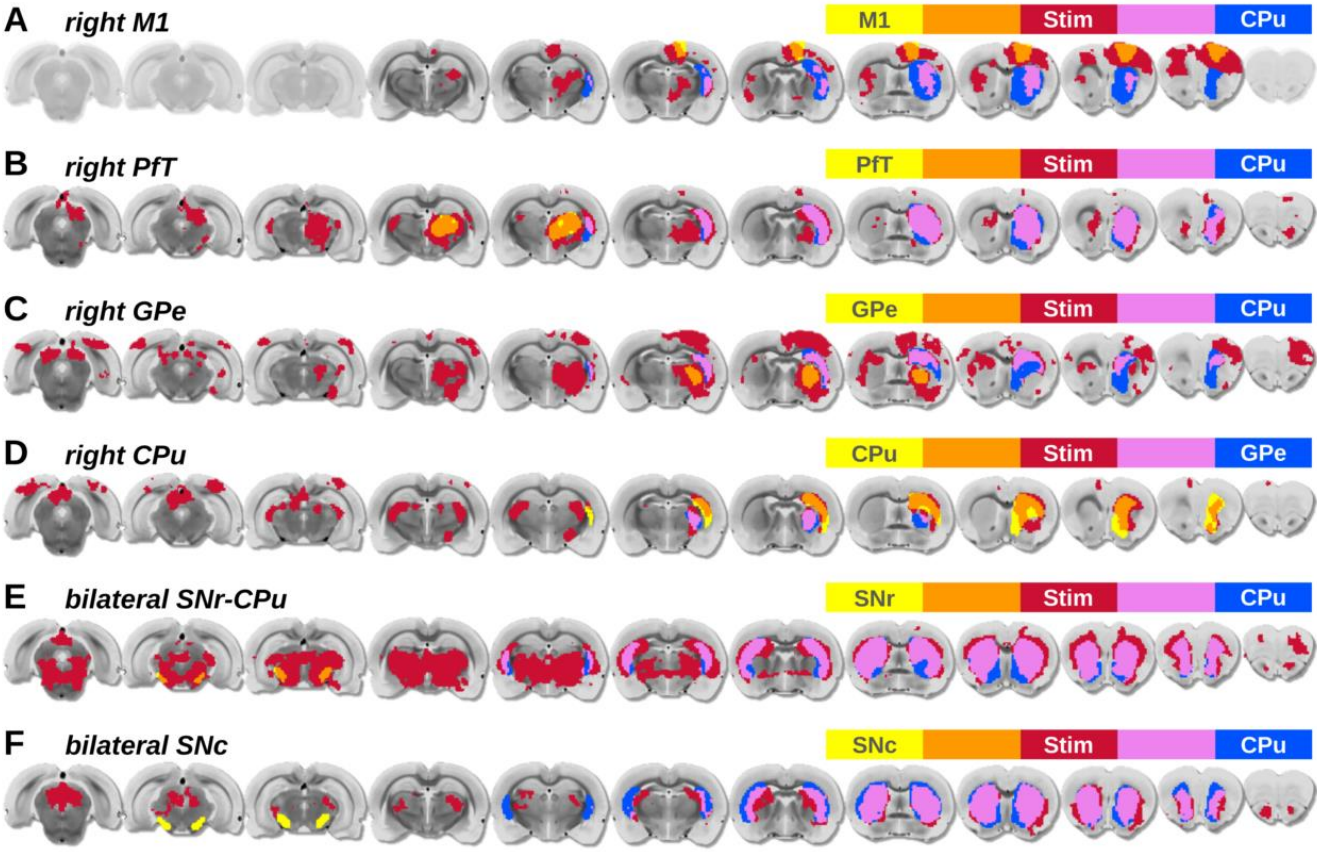


.

**Fig. S3. Extended 12-slice optogenetic fMRI ROI maps from CPu circuit manipulations in Figure 1.** (**A-F**) ROIs used for timeseries extraction were collected from the intersection of the stimulus evoked response maps (red) and anatomical stimulation targets (yellow) or downstream anatomical areas of interest (blue), corresponding to the orange and purple regions, respectively. (**F**) Note, no stimulus evoked response was detected within the anatomical boundaries of SNc for bilateral SNc dopamine neuron stimulation. Left-to-right corresponds to 1 mm steps in the posterior-to-anterior direction, with the 5th slice from the right located at the anterior commissure (approximately –0.36 mm AP).

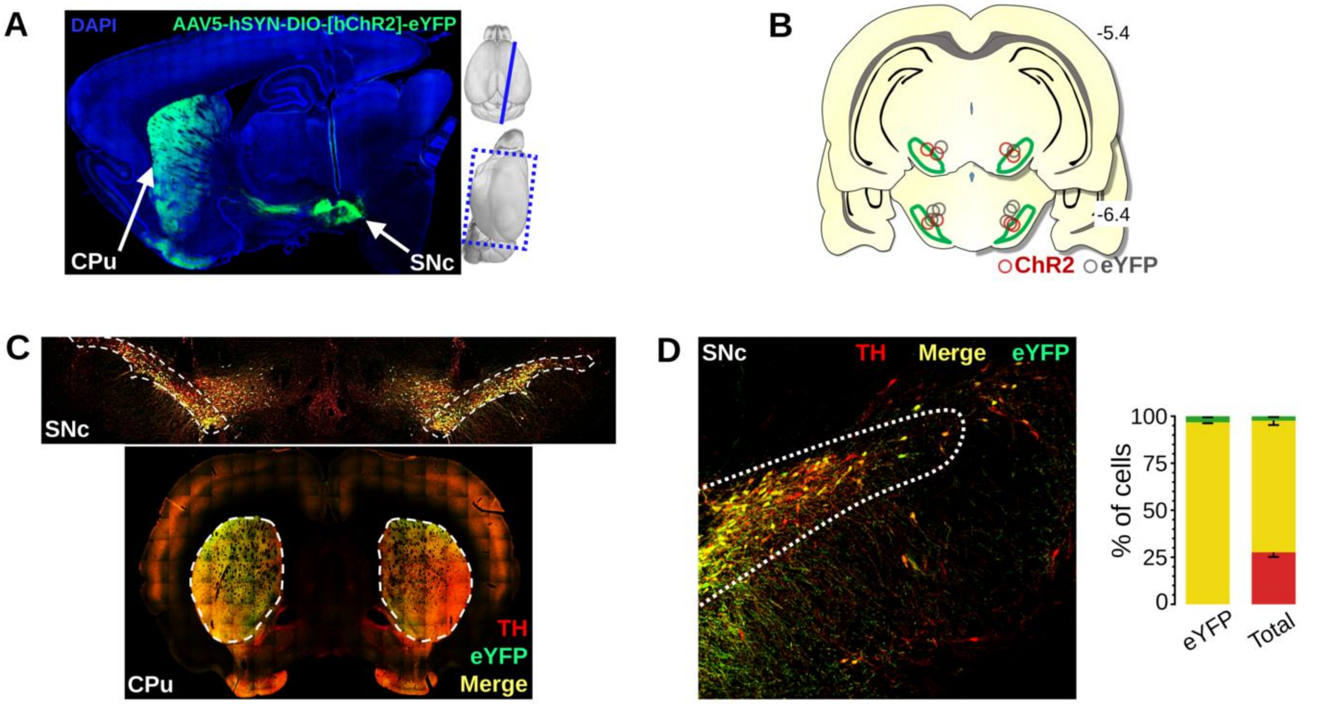


**Fig. S4. Confirmation of optogenetic fMRI SNc dopamine neuron stimulation targets in Figure 1.** (**A**) Representative sagittal cross-section indicating spread of TH-Cre-selective ChR2-eYFP expression from the injection site in SNc to the projection site in CPu. 3-D models of the brain are to the right, indicating the location of the histological slice. (**B**) Locations of optical fiber tips for SNc optogenetic stimulation with fMRI. Anatomical SNc ROI is outlined in green. (**C**) Representative image for high-resolution immunohistochemical verification of bilateral TH and eYFP coexpression in SNc and striatum. (**D**) Immunohistochemical staining used to verify expression of eYFP fluorophore in TH-expressing cells in the SNc (n = 3 brains, 22 slices, 5911 cells); percentages calculated relative to eYFP-positive cells only (left column) or all cells (right column) in SNc target area.

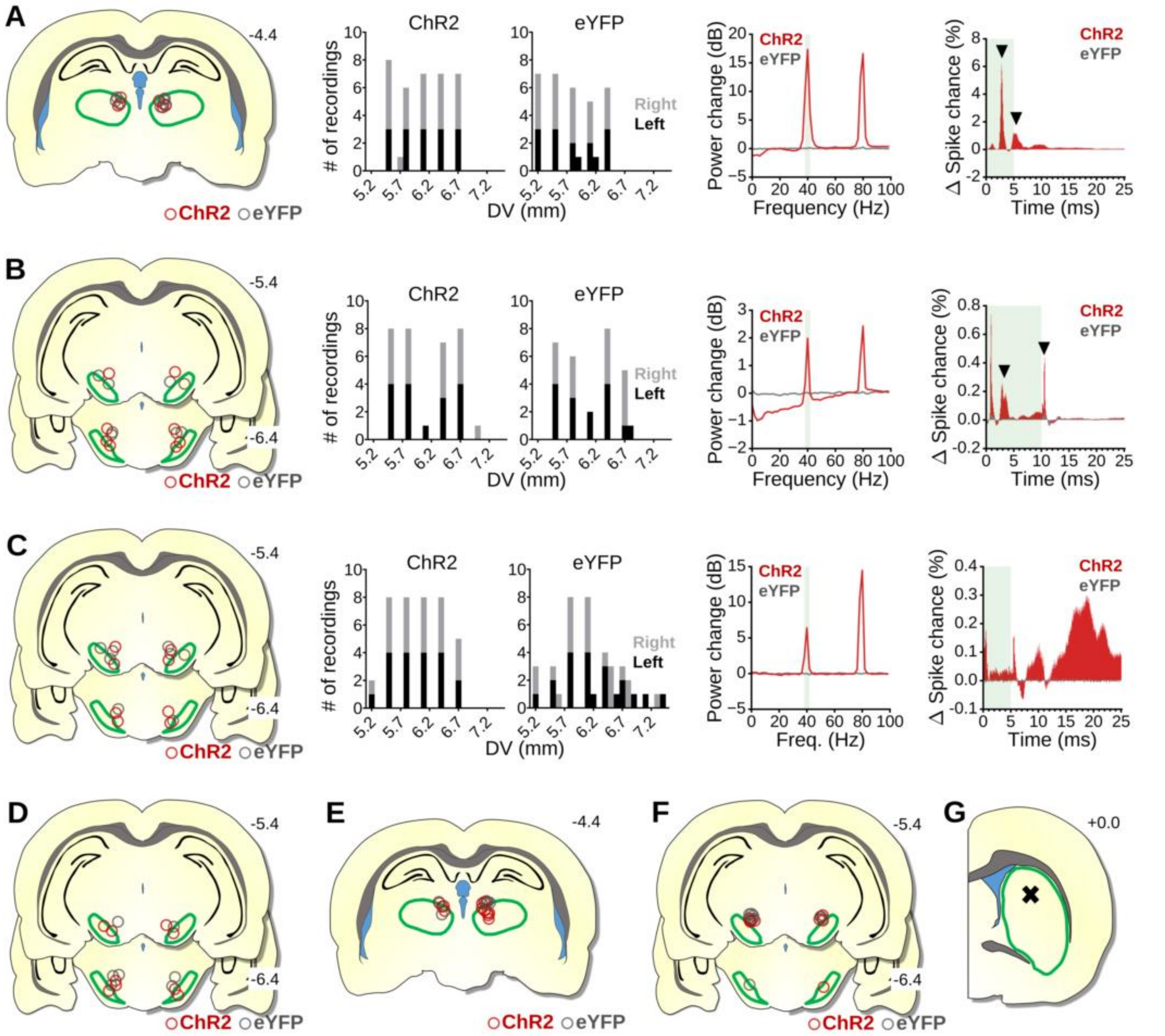


**Fig. S5. Verification of optogenetic stimulation targets, electrode placements, and electrophysiology and FSCV recordings in Figure 2.** (**A-C**) Left to right columns: Locations of optical fiber tips relative to the anatomical stimulation target ROI (outlined in green) for electrophysiology experiments; Acute electrophysiology electrode array implantation depths, where color indicates the recorded hemisphere; Resulting LFP power spectrum change relative to baseline from 40 Hz stimulations; Change in average multi-unit spike probability for the period following each stimulation pulse compared to baseline. (**A**) 10 s, 40 Hz, 5 ms pulse-width PfT stimulation. (**B**) 60 s, 40 Hz, 10 ms pulse-width SNr stimulation. (**A, B,** right column) Black down arrows (left to right) highlight prominent peaks at ~3 ms and at offset of ChR2 stimulation. (**C**) 10 s, 40 Hz, 5 ms pulse-width SNc dopamine neuron stimulation. (**D-F**) From left to right: Locations of optical fiber tips for SNc, PfT, and SNr optogenetic stimulation with FSCV, respectively. (**G**) Acute FSCV electrode implantation depth for both ChR2 and eYFP subjects.

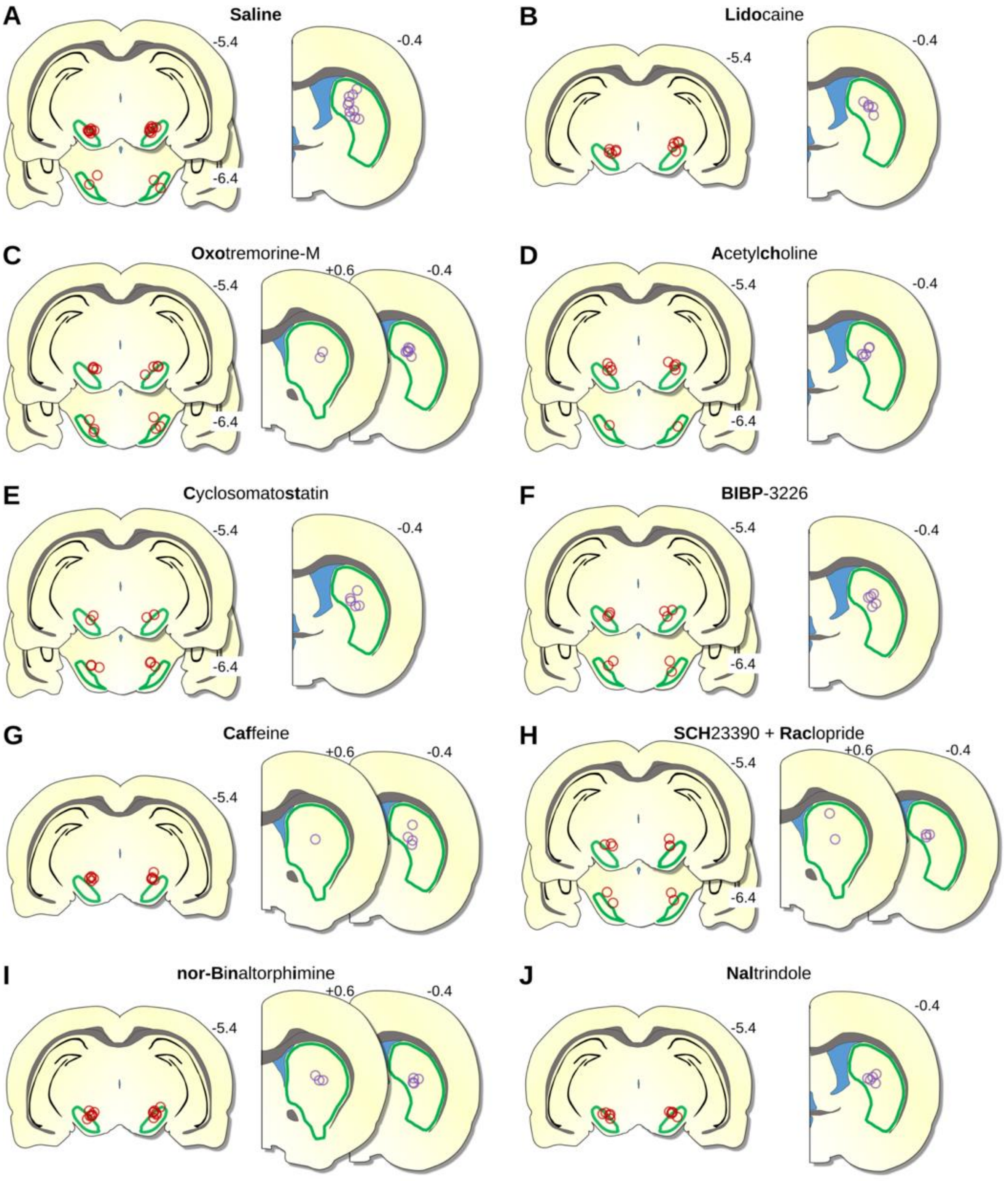


**Fig. S6. Locations of stimulating optical fiber (red) and infusion cannula (purple) tips in subjects from pharmacological CBV-fMRI in Figure 4.** Anatomical boundaries for the optogenetic stimulation site (SNr) and drug infusion site (CPu) are outlined in green.

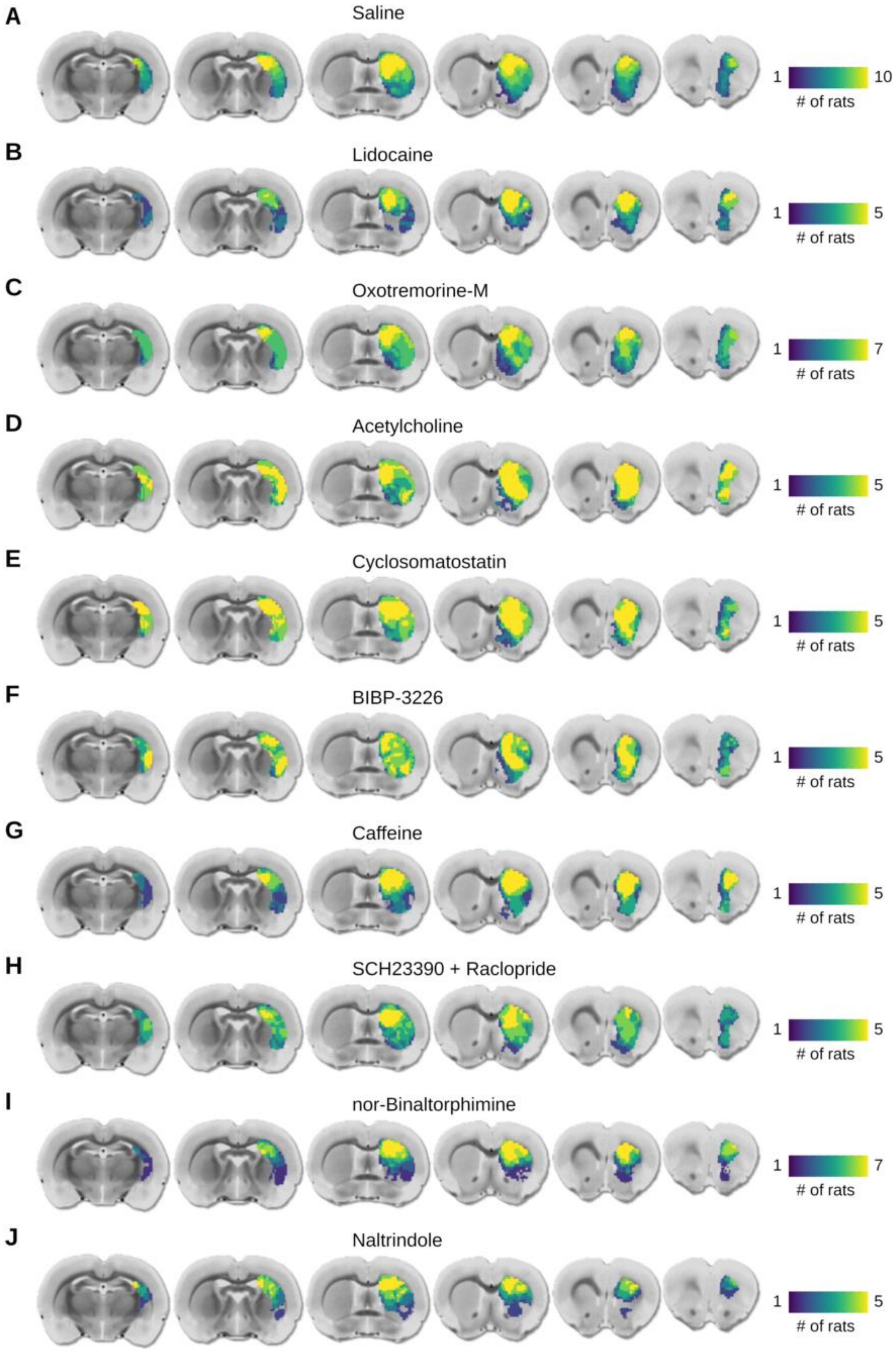
A
Saline
1
10
### of rats
B
Lidocaine
1
5
### of rats
C
Oxotremorine-M
1
7
### of rats
D
Acetylcholine
1
5
### of rats
E
Cyclosomatostatin
1
5
### of rats
F
BIBP-3226
1
5
### of rats
G
Caffeine
1
5
### of rats
H
SCH23390 + Raclopride
1
5
### of rats
I
nor-Binaltorphimine
1
7
### of rats
J
Naltrindole
1
5
### of rats

54

**Fig. S7. Extended 6-slice ROI maps used for pharmacological CBV-fMRI studies in Figure 4.** ROIs used for timeseries extraction were determined by the intersection of individual subject pre-drug evoked response maps and the anatomical boundaries of the right CPu, extending from slices 2-7 out of 12, anterior to posterior. The drug being infused is indicated at the top of each row.

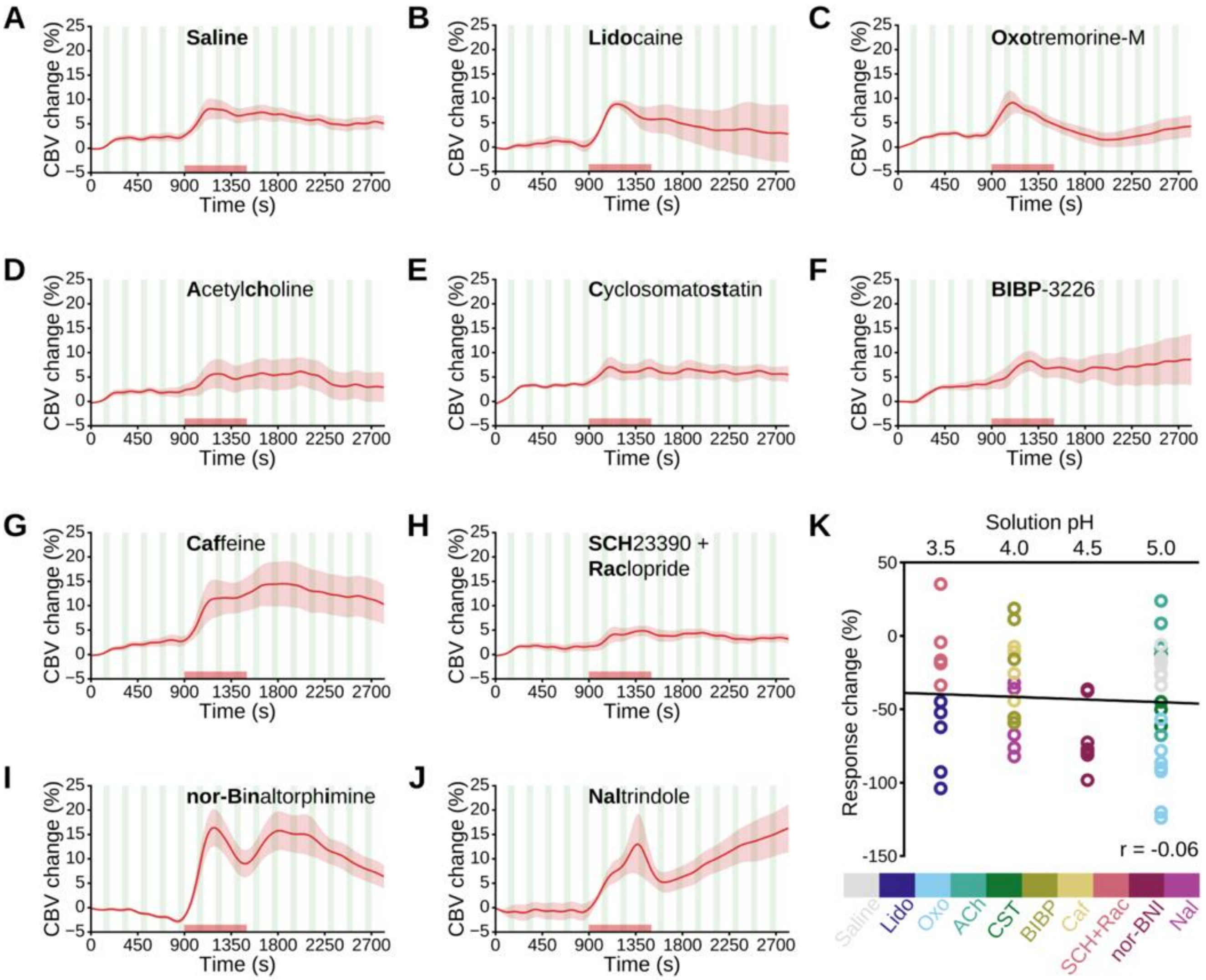


**Fig. S8. Drug infusion controls from pharmacological CBV-fMRI experiments in Figure 4.** (**A-J**) CBV baseline subtraction time-courses as a result of asymmetric least squares and quantile regression methods to correct for drug infusions. The drug being infused is indicated at the top of each panel. Infusion epochs are indicated by a shaded red bar on the x-axis. (**K**) Solution pH values obtained from freshly prepared drug in triplicate, compared to the percent change in CBV baseline during its in vivo infusion. Open circles indicate individual pH readings, color-coded to the appropriate drug name listed below.

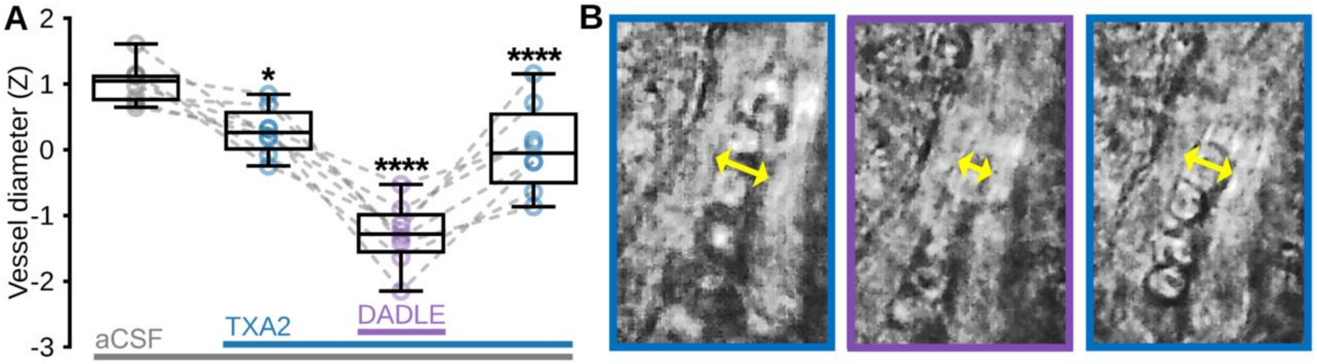


**Fig. S9. Bright-field microscopy reveals acute constriction of CPu microvessels to Enk analog in mouse brain slices.** (**A**) Average microvessel diameter during 5 min experimental epochs (in order: aCSF (baseline), TXA2 + aCSF (pretreatment), DADLE + TXA2 + aCSF (Enk analog), TXA2 + aCSF (washout)) for 8 CPu microvessels (n = 3 mice). Šídák's multiple comparisons test versus preceding experimental condition; *p < 0.05, ****p < 0.0001. (**B**) Representative microvessel showing constriction of vessel diameter (indicated by yellow arrows) to DADLE (purple frame) versus TXA2 pretreatment and washout (left and right blue frames, respectively).

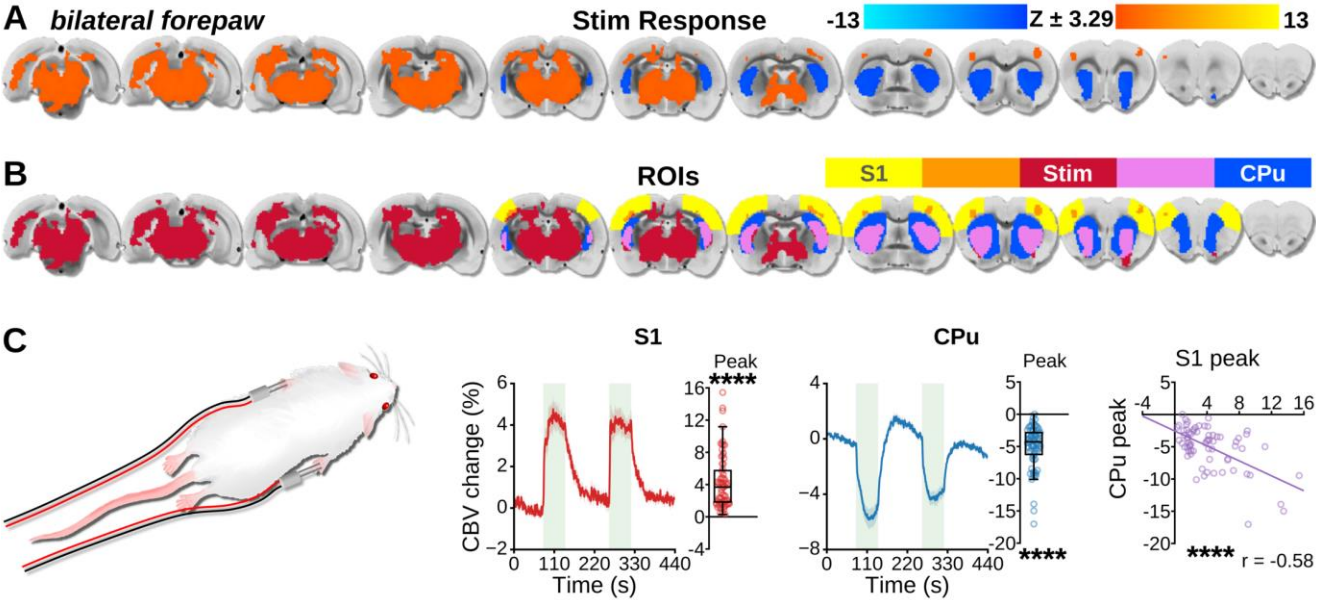


**Fig. S10. Noxious forepaw stimulation elicits negative CBV-fMRI signals in rat CPu.**

Bilateral noxious electrical forepaw stimulation during CBV fMRI produced a significant positive bilateral response in the primary somatosensory cortex (S1) and a robust negative bilateral stimulus-evoked response in CPu (**A**-**C**), and these responses were linearly anticorrelated in peak amplitude (**C**, far right). (**A**) Response maps acquired from subjects following bilateral noxious forepaw stimulation (n = 14 rats, 34 epochs). Response maps thresholded to p<0.001, FWE corrected to α < 0.01. (**B**)

ROIs used for S1 and CPu timeseries extraction were collected from the intersection of the stimulus evoked response maps (red) and anatomical stimulation targets (yellow) or downstream anatomical areas of interest (blue), corresponding to the orange and purple regions, respectively. (**A-B**) Left-to-right corresponds to 1 mm steps in the posterior-to-anterior direction, with the $5^{th}$ slice from the right located at the anterior commissure (approximately –0.36 mm AP). (**C**) Left to right columns: Bilateral forepaw stimulation schematic; CBV time-courses from S1 ROIs, aligned to stimulation epochs ($n = 34$ epochs; green bars indicate optogenetic stimulation blocks; data are presented as mean ±SEM), with corresponding quantified peak amplitude changes ($n = 68$ peaks; one-sample t-test, ****$p < 0.0001$); corresponding CBV time-courses and peak amplitude changes from CPu ROIs; Correlation plots between peak S1 and CPu CBV (linear regression t-test, ****$p < 0.0001$).
